# Cold Shock Fail to Restrain Pre-formed Bacterial Biofilm

**DOI:** 10.1101/324749

**Authors:** Wenying Yu, Qiao Han, Xueying Song, Jiaojiao Fu, Haiquan Liu, Zhuoran Guo, Pradeep K Malakar, Yingjie Pan, Yong Zhao

## Abstract

Environmental temperature fluctuation has great impact on the formation of bacterial biofilm, while little information is available for assessing the influence of sharp temperature shifts on the fate of pre-formed biofilm. In this study, experimental evidence is firstly explored on the response of *Vibrio parahaemolyticus* pre-formed biofilm under cold shock (4 °C and 10 °C). Surprisingly, biofilm biomass of *V. parahaemolyticus* significantly increased during the period of cold shock as revealed by crystal violet staining. Polysaccharides and proteins contents in extracellular polymeric substances were gradually enhanced after cold shocks and exhibited high consistency. RT-qPCR demonstrated the expression of flagella and virulence-related genes were up-regulated. Most of QS and T3SS genes were slightly up-regulated, and three T3SS genes (*vcrD1*, *vcrD2β* and *vopD1*) were down-regulated. Furthermore, the biofilm structure of *V parahaemolyticus* have been analyzed by Confocal laser scanning microscopy (CLSM), which sharply changed under cold shocks. The correlation analysis further displayed the significant correlation (P < 0.01) among biofilm structure parameters, and weak correlation (P < 0.05) between biofilm related genes and biofilm structure parameters. In conclusion, our results novel discovered that *V. parahaemolyticus* biofilm related genes were actively expressed and biofilm biomass was continuously increased, biofilm structure was tremendously changed after cold shock. This study underscored the risk that biofilm cells had the ability to adapt to low temperature shift.

**IMPORTANCE:** Biofilms are widespread in natural environments, especially on the surface of food and medical biomaterials, which threaten human safety from persistent infections. Previous studies simply focused on biofilm formation of microorganisms under steady state, however, the actual environment frequently fluctuated. *V. parahaemolyticus* is a widely distributed foodborne pathogen, temperature play a great role in its survival. Researchers generally assume that cold environment can restrain biofilm formation and bacterial activity. This study explored the effects of *V. parahaemolyticus* biofilm upon a shift from 37 °C to 4 °C or 10 °C from two aspects. On the one hand, the changes of biofilm biomass and EPS contents, the expression of biofilm related genes directly described that pre-formed bacterial biofilm could not be controlled efficiently in cold environment. On the other hand, the CLSM images revealed biofilm morphological structure change, the correlation analysis showed inner relationship among biofilm structure parameters and biofilm related genes. These results suggested that cold shock fail to restrain pre-formed bacterial biofilm, therefore be a potential risk in nature environment.

## 1. Introduction

A biofilm is a multicellular complex, formed of microorganisms that are attached to a surface and generally embedded in extracellular polymeric substances (EPS) (1,2,3). Polysaccharides and proteins are the most prevalent components of EPS in biofilms, the production of mature biofilms requires polysaccharides to hold the cells together and maintains the structure integrity of the biofilm (4,5). Biofilms is a means of persistence for bacteria which play a crucial role for bacterial life cycle (6). According to Center for Disease Control and Prevention (CDC), almost 65 % of all reported bacterial infections are caused by bacterial biofilms (7). Bacterial cells in biofilm are more resistant towards disinfectants and antibiotics than their planktonic forms (8,9). The capacity of bacterial biofilm formation can be directly affected by environmental temperature (10,11). Additionally, temperature also affects biofilm formation by modulates bis-(3’-5’)-cyclic dimeric guanosine monophosphate (c-di-GMP) levels in Gram-negative bacterial (12,13). As bacterial second messengers, c-di-GMP involved in several cellular processes which including virulence, biofilm formation, and flagellar synthesis, and so on (14). For the environmental temperature fluctuated quickly, the phenomenon of pre-formed biofilm exposed to cold shock is a well recognized but neglected phenomenon which has been discussed in this article.

*V. parahaemolyticus* is a halophilic, Gram-negative, curved rod-shaped bacterium (15), widely distributed in aquatic reservoirs (16,17), could persist in food processing environments by forming biofilm (18). *V. parahaemolyticus* has been considered as a leading cause of seafood-derived illness in many Asian countries, including China, Japan, and Korea (19,20,21). *V. parahaemolyticus* outbreaks are highly correlated with temperature fluctuation (22,23).

According to the research of Cook et al., the growth of *V. parahaemolyticus* at planktonic form was inhibited when temperature was below 10 °C (24,25). However, Costerton et al. reported that the bacteria in biofilm are more resistant to low temperatures than free counterparts (1). Han et al. found that *V. parahaemolyticus* at 4 °C and 10 °C also could form biofilm as monolayers, but the biofilm biomass was significantly decreased (*p* < 0.05) compared with higher temperatures (15 °C - 37 °C) (26). Specially for the transport,retail and processing of commercial seafoods which are typically stored at 4 °C to 10 °C (27,28) and food-borne pathogens in seafoods encounter significant temperature fluctuation.

Bacterial quorum sensing (QS), flagella, type III secretion systems (T3SS) and two haemolysins (TDH and TRH) can regulate the biofilm formation of *V. parahaemolyticus* through the expression of the related genes. Bacterial QS is a communication process among cells and could monitor cell population density through auto-inducers (29), which is essential for bacterial biofilm formation (30,31). AphA is an activator of virulence and biofilm formation in *V. parahaemolyticus*, OpaR represses biofilm formation in the pandemic O3:K6 *V. parahaemolyticus* (32). The expression of *pilA* (chitin-regulated pilus pilin gene) led to increase the formation of the biofilms by chitin. In addition, two type III secretion systems (T3SS1 and T3SS2), the thermostable direct haemolysin (TDH) and the TDH-related haemolysin (TRH) made main contributions to the pathogenicity of *V. parahaemolyticus* (33). T3SS1 and T3SS2 genes have relation with the biofilm production in *V. parahaemolyticus* (34).

To date, little information is available on pre-formed bacterial biofilm changes at low-temperature shifts, although it is crucial for environmental protection and controlling bacterial infections outbreaks. Thus, in this study, we investigated the fate of *V. parahaemolyticus* biofilm upon a shift from 37 °C to 4 °C or 10 °C, and analyzed the changes of biofilm biomass, viable cells, EPS contents, biofilm architecture and the expression of biofilm related genes.

## 2. RESULTS

### 2.1 Biofilm biomass changes

The biomass changes of biofilm (A) and dynamic changes of viable cells in biofilm forms (B) under cold shock conditions were illustrated in Figure 1. Overall, with the growth of incubation time, biofilm biomass was increased (Figure 1A); the viable cells in planktonic were gradually decreased with the increasing of treatment time (Figure 1B). Under cold environmental (4 °C and 10 °C), the OD_570nm_ of pre-form biofilm was 0.399 which keeping growth within 60h; the viable cells of *V. parahaemolyticus* declined from 8.25 Log_10_ CFU/ml to 7.35 Log_10_ CFU/ml, 6.52 Log_10_ CFU/ml, respectively. In addition, there was no significant difference (*p*>0.05) between the viable cells in *V. parahaemolyticus* biofilm exposed to 4 °C and 10 °C.

**Figure 1.**
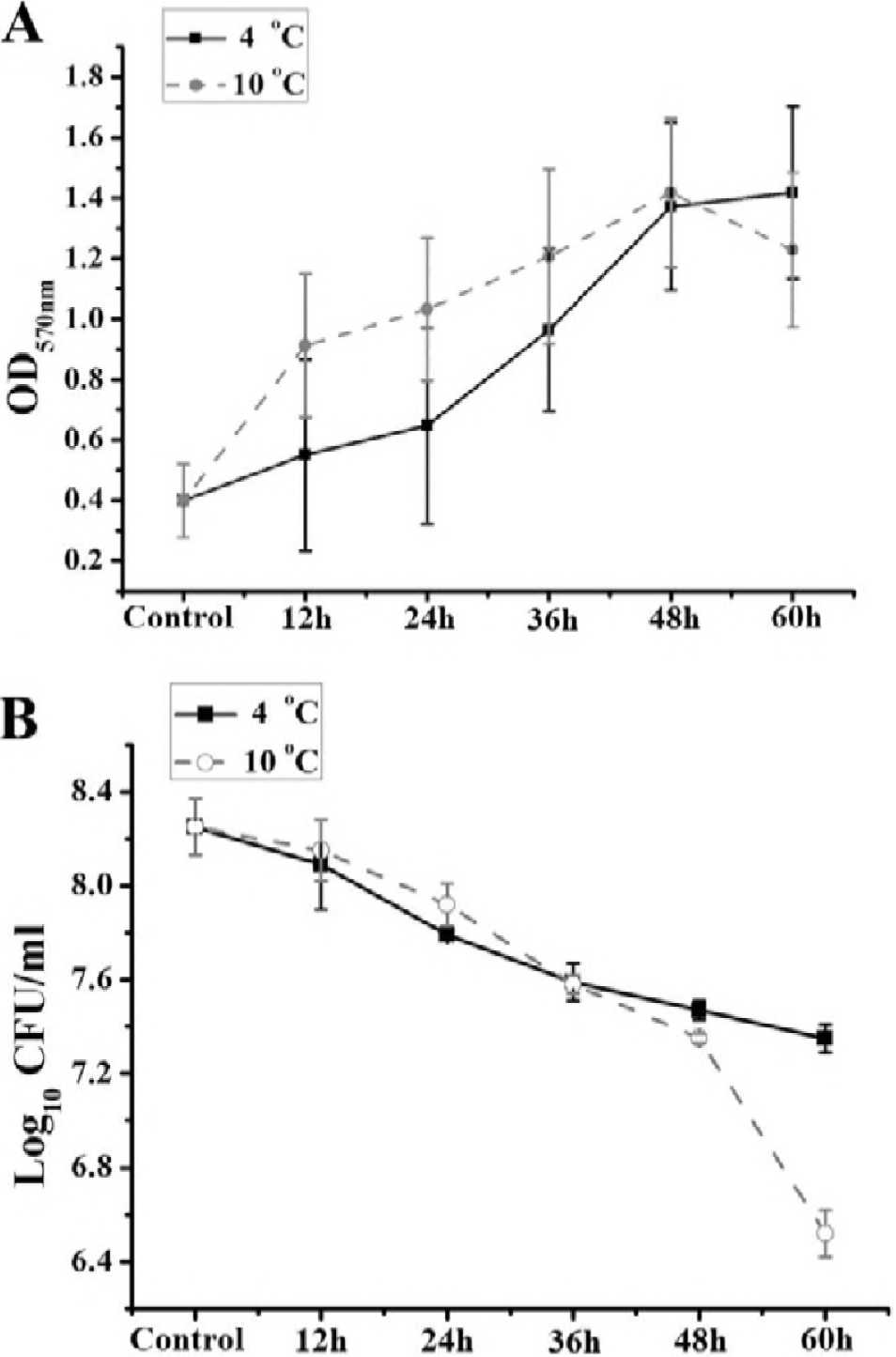
The change of biofilm biomass (A) of *V. parahaemolyticus* exposed to cold shock (4 °C and 10 °C). The viable biofilm bacteria (B) were counted after cold shock (4 °C and 10 °C). The data are presented as the mean of log values of the bacterial population ± standard deviation for three independent replicates.

### 2.2 EPS analysis

To evaluate the effects of cold shock on EPS production, we analyzed the total exopolysaccharides and proteins in EPS of the pre-formed biofilm treated from 12 h to 60 h. As shown in Figure 2, the exopolysaccharides contents increased continuously, and no remarkable difference of exopolysaccharides contents between 4 °C and 10 °C. The results demonstrated that cold shock could only reduce the growth rate of exopolysaccharides, rather than restrain it absolutely, which coincident with the results from the crystal violet staining. Meanwhile, Figure 2C analyzed the linear correlation between protein and polysaccharides in biofilm. The linear fitting results were list as followed.

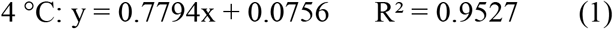

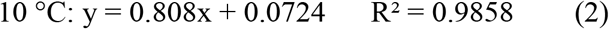

**Figure 2.**
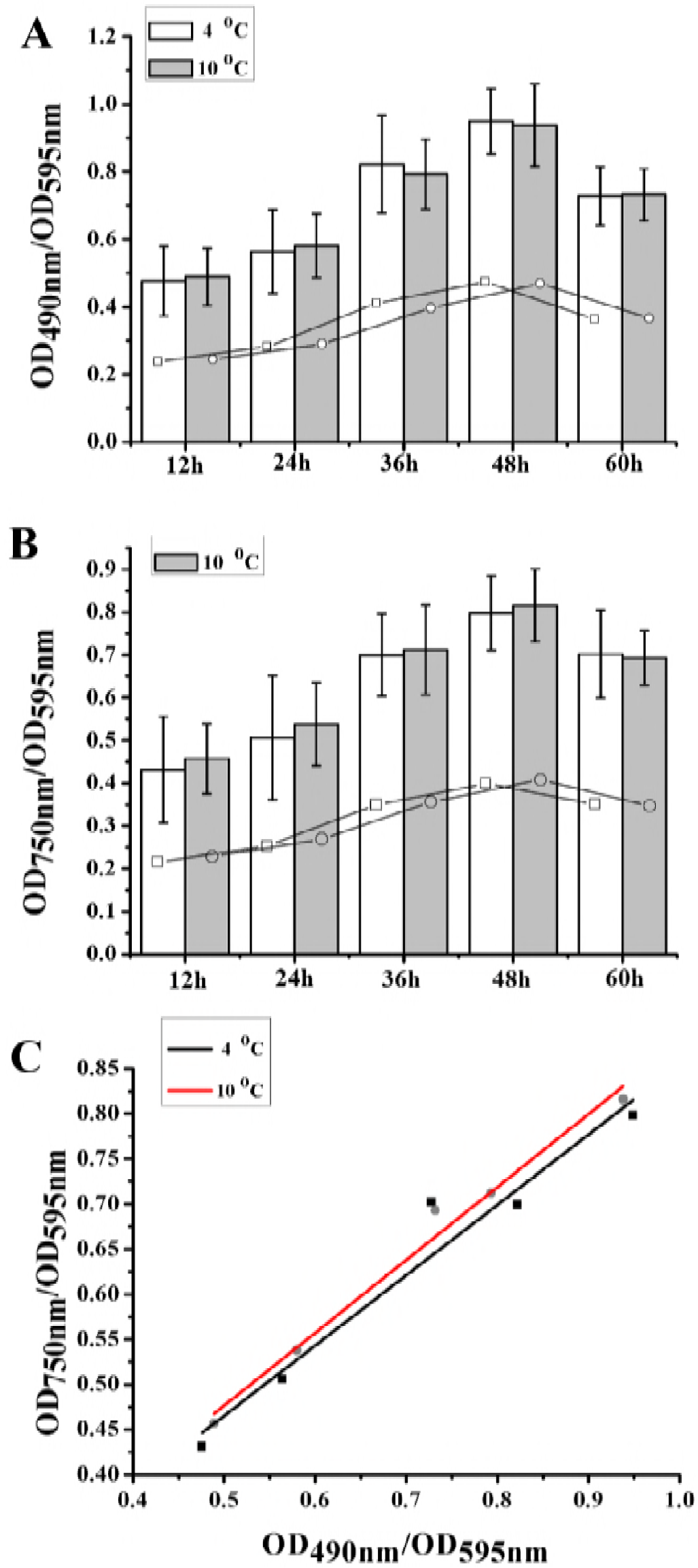
The contents change of EPS constituent exposed to cold shock (4 °C and 10 °C). (A) Total contents (column) of exopolysaccharides in EPS of *V. parahaemolyticus* biofilm, and the average value (line) which showed the variation tendency of exopolysaccharides after cold shock. (B) Total contents (column) of proteins in EPS of *V. parahaemolyticus* biofilm, and the average value (line) which showed the variation tendency of proteins after cold shock. Error bars represent the standard deviations of 3 measurements. * indicates significant difference (p < 0.05). (C) The linear regression of total contents of exopolysaccharides and proteins in EPS of *V. parahaemolyticus* biofilm.

The higher R^2^ indicates a higher degree of linear dependence and the consistency of protein and polysaccharides. Meanwhile, the pearson correlation coefficients were calculated that 0.976 and 0.993 at 4 °C and 10 °C, respectively. The correlations of protein and polysaccharides at two temperature conditions were both significant at the 0.01 level (2-tailed). Those results indicated that the compositions of EPS are high consistency and the order of biofilm structure are stronger at cold environment with the increase of biofilm biomass.

### 2.3 Gene expression analysis

In order to better understand the fate of pre-formed biofilm of *V. parahaemolyticus* exposed to cold shock, we further analyzed the biofilm-related gene expression changes, including flagella (*pilA*), QS (*aphA*, *opaR*), virulence (*trh*) and T3SS (*vcrDl*, *vopS*, *vopD1*, *vscC2β*, *vcrD2β*, *vopP2β*). As shown in Figure 3, all of the selected flagella and virulence genes were significantly up regulated in the biofilm cells. Also, with the increase of incubation time, the genes *aphA* and *vscC2β* and *vopP2β* were up regulated gradually, whereas the genes *opaR* and *vopS* were expressed without significant difference. However, after cold shock, T3SS genes (*vcrD1, vcrD2β* and *vopD1*) were down-regulated. Earlier studies showed that *vcrD1* and *vcrD2β* encoding an inner membrane protein (35), and *vopD1* are essential for translocation of T3SS1 effector of *V. parahaemolyticus* (36).

**Figure 3.** Expression profiles of a selected set of genes in *V. parahaemolyticus* biofilm exposed to 4 °C and 10 °C for 12h, 24h, 36h, 48h and 60h. Induced expression is represented in yellow, repressed expression is represented in blue, and little changed expression is represented in black. Differential expression of genes involved in flagella, QS, virulence gene, T3SS1 and T3SS2 was observed upon a cold shift. The color scale is shown at the lower right corner. Primers sequences used in RT-qPCR assay are provided in Table 1.

**Table 1.**
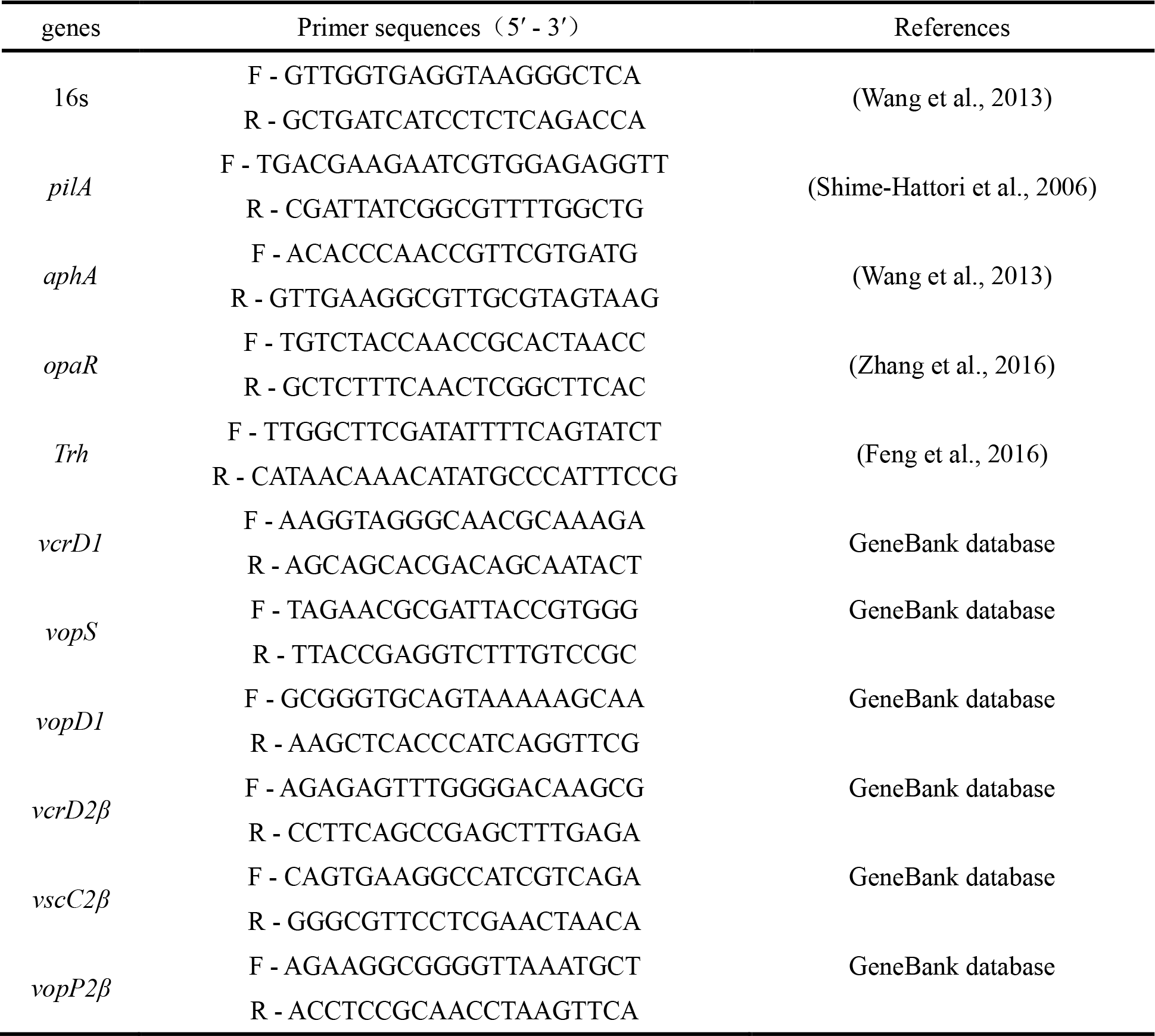
Primer sequences of the RT-qPCR assay.

The results showed that the biofilm cells have adapted to the low temperature shift and form a more stable structure to defend the adverse environment. This was consistent with the findings of biofilm biomass changes and EPS changes.

### 2.4 Biofilm architecture changes

The CLSM images of pre-formed *V. parahaemolyticus* biofilm were presented in Figure 4, showing that *V. parahaemolyticus* had the ability to form complex three-dimensional structures. Figure 5 indicated that, with the increasing of incubation time, biofilm architecture ranged from a flat homogeneous layer of cells to a complex structure. Specially, the contact surfaces were completely covered by dense and homogeneous biofilm when cultured at 4 °C and 10 °C for 36 h.

**Figure 4.**
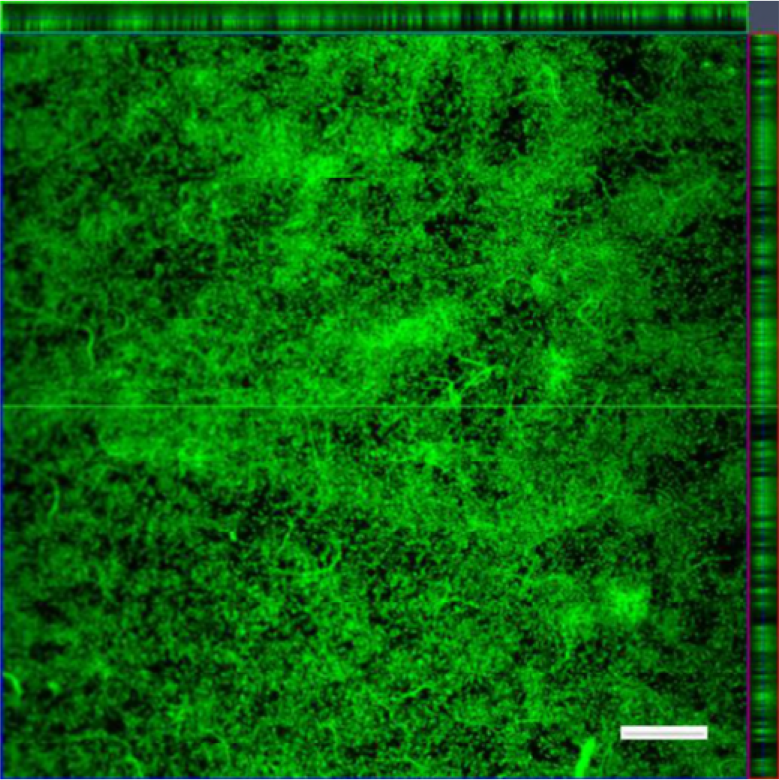
Confocal laser scanning microscopy (CLSM) images of pre-formed biofilm formed by *V. parahaemolyticus* at 37 °C for 24 h. The scale bar represents 20 μm.

**Figure 5.**
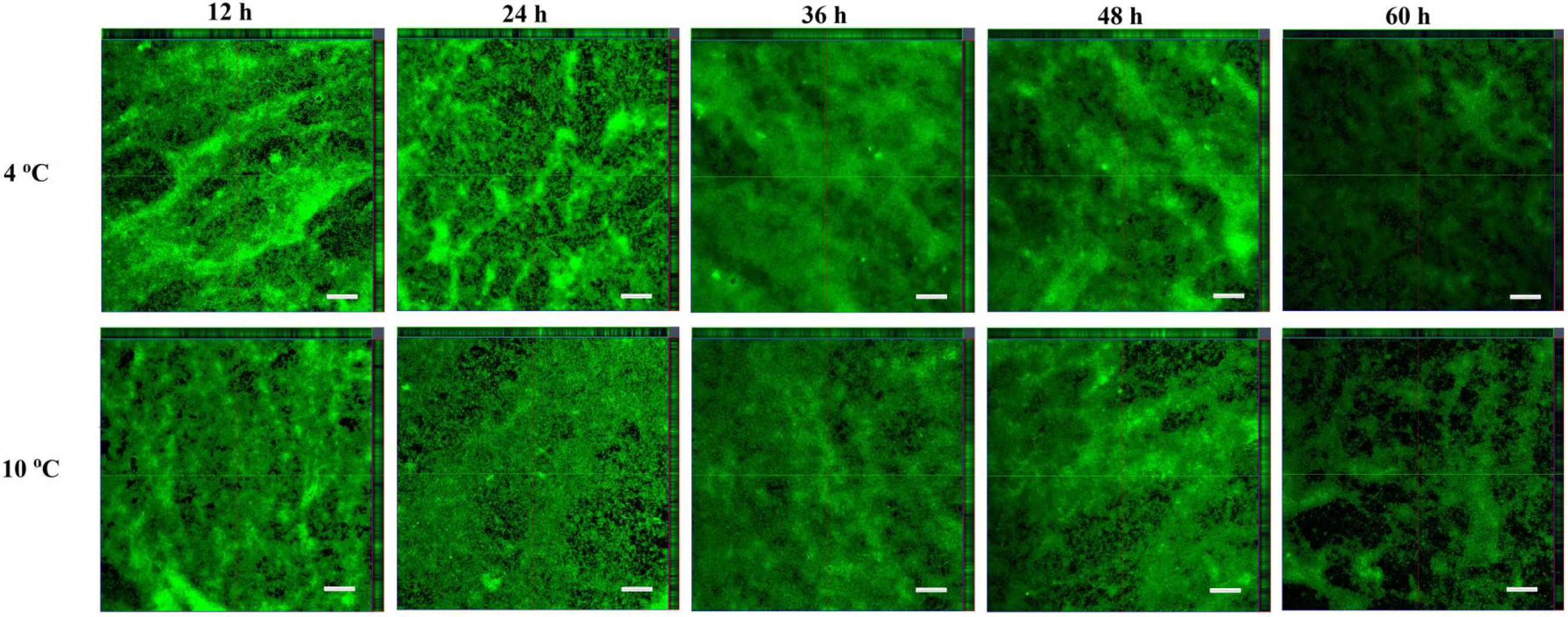
Confocal laser scanning microscopy (CLSM) images of biofilm formed by *V. parahaemolyticus* subjected to cold shock (4 °C and 10 °C). The biofilms were incubated at 37 °C for 24 h to obtain preformed biofilm, and then shifted to 4 °C and 10 °C immediately or kept at 37 °C for 12 h, 24 h, 36 h, 48 h and 60 h. The scale bar represents 20 μm.

Quantitative parameters describing biofilm physical structure were summarized in Figure 6. The MT provides a measure of the spatial size of the biofilm and is the most common variable used in biofilm literature. The MT value of pre-formed biofilm was 1.77 and slightly changed when exposed to cold shock at 10 °C, while it was tremendously changed at 4 °C. Average diffusion distances (ADD) have been suggested as measurement of the distance over that nutrients and other substrate components diffused from the voids to the bacteria within micro-colonies (37,38). The ADD value of pre-formed biofilm was 1.04. The ADD of biofilm at the low temperatures (4 and 10 °C) were higher than the initial value within 36 h.

**Figure 6.**
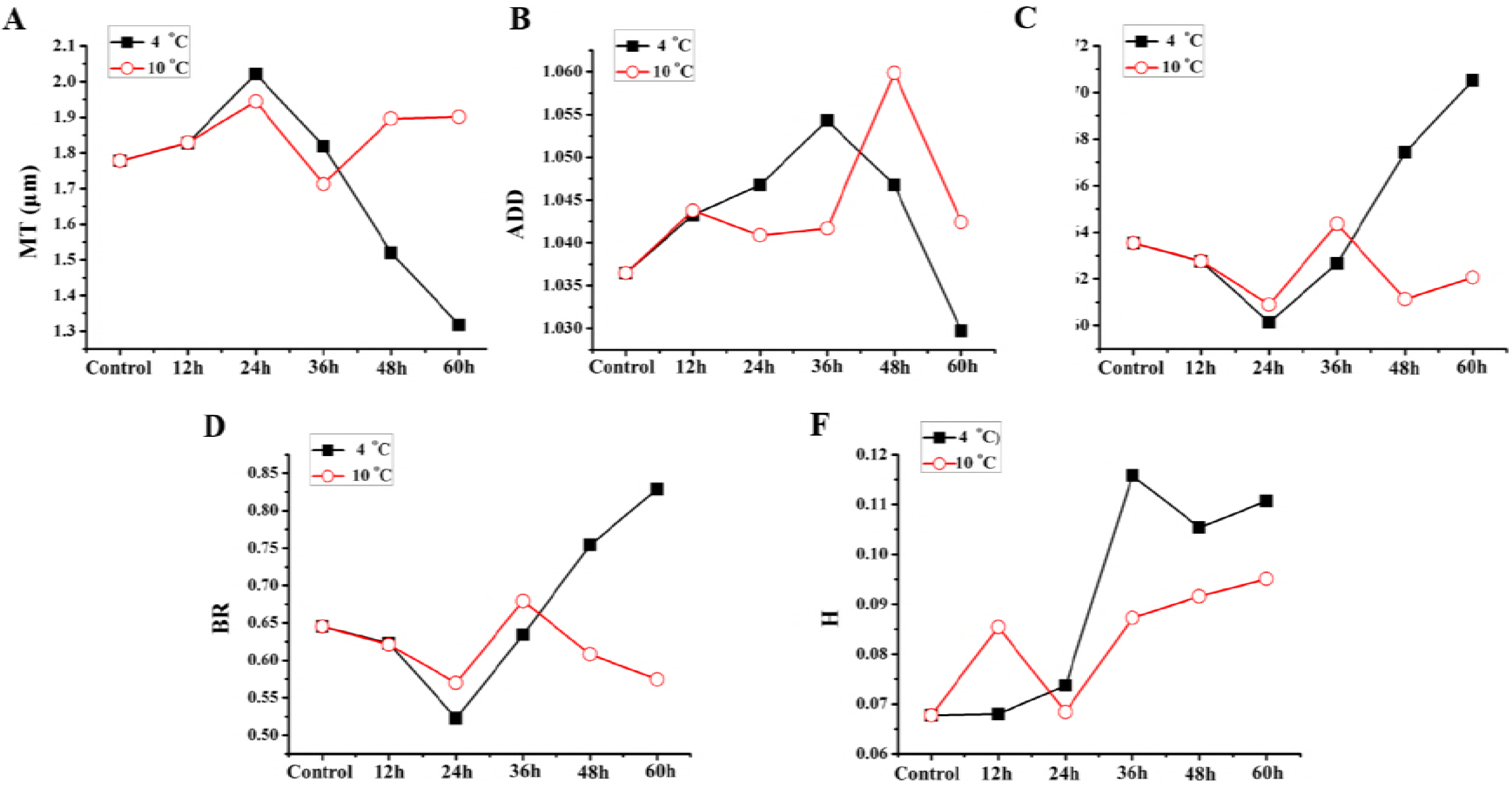
Quantification of structural parameters changes (3A-F) in biofilm formed by *V. parahaemolyticus* after low temperature shift, including mean thickness (MT), average diffusion distance (ADD), porosity (P), biofilm roughness (BR) and homogeneity (H).

In addition, the biofilm roughness (BR) indicates the variability in the biofilm thickness but does not describe the variability in the surface of the cell clusters (39). The BR value of pre-formed biofilm was 0.645295. After cold shock (4 °C), the BR value was significantly increased (Figure 6E). Areal parameters describe the morphological structures of biofilm. We selected porosity (P) as a textural parameter. The porosity decreased with increasing number of cell clusters. The porosity value of pre-formed biofilm was 0.64. Figure 6D showed that after cold shock (4 °C), the porosity value was significantly increased, in the range from 0.601 to 0.705, with the increasing of incubation time. Textural entropy is a measure of randomness in the gray scale of the image. The higher the textural entropy, the more heterogeneous the image is. The homogeneity value of pre-formed biofilm was 0.059572.

When shifted to low temperatures, the pre-formed biofilm needed to accommodate the changed circumstances to form mature biofilm. The CLSM images were in accordance with the results of crystal violet staining. These results indicated that cold shock could only prolonged the period of biofilm mature.

### 2.5 Correlation analysis

The correlation among biofilm structure parameters were list in Table2. Correlation is significant at the 0.01 level (2-tailed). Analysis of 6 different biofilm structure parameters of *V. parahaemolyticus* in two temperature conditions showed a novel positive correlation between biofilm thickness (BR) and porosity (P). The negative correlation between spatial size of the biofilm (MT) and biofilm thickness (BR); spatial size of the biofilm (MT) and porosity (P). Compared with 10 °C, those trends are more obvious at 4 °C.

**Table 2.** The correlation among biofilm structural parameters.

| Temperature<br>(°C) | Biofilm<br>structural<br>parameters | MT |  | BR |  | P |  | ADD |  | H |  |
| --- | --- | --- | --- | --- | --- | --- | --- | --- | --- | --- | --- |
|  |  | A | B | A | B | A | B | A | B | A | B |
| 4 | MT | 0 | 1 | 0.000 | -0.996** | 0.000 | -0.999** | 0.196 | 0.692 | 0.204 | -0.683 |
|  | BR |  |  | 0 | 1 | 0.001 | 0.992** | 0.243 | -0.642 | 0.182 | 0.707 |
|  | P |  |  |  |  | 0 | 1 | 0.177 | -0.713 | 0.222 | 0.664 |
|  | ADD |  |  |  |  |  |  | 0 | 1 | 0.863 | -0.108 |
|  | H |  |  |  |  |  |  |  |  | 0 | 1 |
| 10 | MT | 0 | 1 | 0.007 | -0.968** | 0.008 | -0.966** | 0.729 | 0.215 | 0.640 | -0.287 |
|  | BR |  |  | 0 | 1 | 0.055 | 0.870 | 0.987 | -0.01 | 0.697 | 0.240 |
|  | P |  |  |  |  | 0 | 1 | 0.497 | -0.406 | 0.600 | 0.320 |
|  | ADD |  |  |  |  |  |  | 0 | 1 | 0.540 | 0.370 |
|  | H |  |  |  |  |  |  |  |  | 0 | 1 |

Note: A, Sig. (2-tailed); B, Correlation coefficient; \*, correlation is significant at the 0.05 level (2-tailed); \*\*, correlation is significant at the 0.01 level (2-tailed).

In the present study, Table3 displayed the correlation between biofilm formation related genes and biofilm structural parameters at 4 °C and 10 °C. The significant correlations were exhibited between *pilA* and H; *vopP2β* and MT, BR at the 0.05 level (2-tailed); *vcrD1* and MT, BR, P; *vscC2β* and ADD. In conclusion, after cold shock, the flagella and T3SS genes have significant correlation with biofilm structural in *V. parahaemolyticus*.

**Table 3.** The correlation between biofilm structural parameters and biofilm formation related genes.

| Genes | Tempe-<br>rature<br>(°C) | MT |  | BR |  | P |  | ADD |  | H |  |
| --- | --- | --- | --- | --- | --- | --- | --- | --- | --- | --- | --- |
|  |  | A | B | A | B | A | B | A | B | A | B |
| *pilA* | 4 | 0.178 | -0.711 | 0.168 | 0.722 | 0.186 | 0.702 | 0.835 | -0.13 | **0.013** | **0.951\*** |
|  | 10 | 0.773 | 0.179 | 0.818 | -0.143 | 0.740 | -0.205 | 0.293 | 0.592 | **0.019** | **0.936\*** |
| *aphA* | 4 | 0.494 | 0.409 | 0.458 | -0.44 | 0.513 | -0.393 | 0.636 | -0.290 | 0.068 | -0.850 |
|  | 10 | 0.804 | 0.155 | 0.629 | -0.296 | 0.989 | 0.009 | 0.402 | -0.490 | 0.797 | -0.160 |
| *opaR* | 4 | 0.636 | -0.290 | 0.719 | 0.223 | 0.589 | 0.329 | 0.170 | 0.720 | 0.518 | -0.388 |
|  | 10 | 0.902 | 0.077 | 0.958 | 0.033 | 0.762 | -0.188 | 0.165 | 0.726 | 0.293 | 0.592 |
| *Trh* | 4 | 0.565 | 0.349 | 0.573 | -0.342 | 0.566 | -0.348 | 0.887 | -0.089 | 0.111 | -0.790 |
|  | 10 | 0.936 | 0.05 | 0.881 | -0.094 | 0.992 | 0.006 | 0.622 | -0.301 | 0.535 | -0.374 |
| *vcrD1* | 4 | **0.021** | **-0.932\*** | **0.017** | **0.941\*** | **0.026** | **0.921\*** | 0.224 | -0.661 | 0.165 | 0.725 |
|  | 10 | 0.728 | 0.215 | 0.782 | -0.172 | 0.682 | -0.253 | 0.559 | 0.354 | 0.578 | 0.338 |
| *vcrD2β* | 4 | 0.450 | 0.448 | 0.536 | -0.373 | 0.411 | -0.482 | 0.172 | 0.718 | 0.776 | -0.177 |
|  | 10 | 0.240 | -0.645 | 0.148 | 0.746 | 0.395 | 0.497 | 0.376 | 0.514 | 0.288 | 0.597 |
| *vopS* | 4 | 0.342 | -0.545 | 0.371 | 0.519 | 0.330 | 0.557 | 0.744 | -0.202 | 0.180 | 0.709 |
|  | 10 | 0.175 | -0.715 | 0.120 | 0.780 | 0.280 | 0.605 | 0.444 | 0.452 | 0.225 | 0.660 |
| *vopD1* | 4 | 0.755 | -0.194 | 0.695 | 0.242 | 0.779 | 0.174 | 0.341 | 0.546 | 0.294 | 0.591 |
|  | 10 | 0.554 | 0.358 | 0.521 | -0.386 | 0.606 | -0.315 | 0.940 | 0.047 | 0.778 | 0.175 |
| *vopP2β* | 4 | 0.459 | 0.436 | 0.513 | -0.393 | 0.394 | -0.497 | 0.138 | 0.758 | 0.797 | 0.160 |
|  | 10 | **0.033** | **-0.907\*** | **0.010** | **0.958\*** | 0.106 | 0.797 | 0.872 | -0.101 | 0.676 | 0.257 |
| *vscC2β* | 4 | 0.516 | 0.390 | 0.606 | -0.315 | 0.470 | -0.430 | 0.098 | 0.808 | 0.779 | 0.175 |
|  | 10 | 0.994 | 0.004 | 0.773 | 0.179 | 0.761 | -0.189 | **0.008** | **0.966\*\*** | 0.346 | 0.541 |

Note: A, Sig. (2-tailed); B, Correlation coefficient; \*, correlation is significant at the 0.05 level (2-tailed); \*\*, correlation is significant at the 0.01 level (2-tailed).

## 3. DISCUSSION

The presence of biofilms could protect cells from chemical sanitizers and environment stress, so the biofilm-forming capability of *V. parahaemolyticus* allows its persistence and transmission in environment (40). *V. parahaemolyticus* is a serious threat to public health. Previous studies focused on the biofilm formation at various constant temperatures (11,41), while the actual environmental were fluctuated frequently. In the present study, we gained further insights into the fate of pre-formed *V*. *parahaemolyticus* biofilm under cold shock.

A striking finding was that some survival biofilm cells of *V. parahaemolyticus* gradually adapted to cold shock and protected themselves against the cold shift. Figure 1 showed that the viable bacteria counts in pre-formed biofilm were declined, while biofilm biomass were increased when shifted to low temperature (4 °C and 10 °C). The presence of this phenomenon may depend on the viable but non-culturable (VBNC) state of *V. parahaemolyticus* at low temperature shift. Previous research revealed that under the stressed environments, *V*. *parahaemolyticus* enters into self-defensive mode of metabolism or a physiologically VBNC state (42).

After cold shock, the genes related with flagella, virulence and QS (*aphA*) were up-regulated which suggesting there is a protective effect on the survival biofilm cells. Besides, cold shock could also have a deep influence on biofilm morphological structure. As it showed in Table3, flagella and T3SS genes (*pilA*, *vcrD1*, *vopP2β* and *vscC2β*) have significant correlation with the structural parameters which may regulate the biofilm structure under cold shift in *V. parahaemolyticus*. It has been reported that flagella are involved in the biofilm formation, especially for *Vibrio spp*., usually by enhancing movement towards the surface of biofilm (5,43). Enos-Berlage et al. tested biofilm formation of *V. parahaemolyticus* and observed that *flgE* and *flgD* mutants were defective in attachment and biofilm formation (44). Asadishad et al. also found that bacterial swimming motility was decreased and the transcription of flagellin encoding genes were expressed after *Bacillus subtilis* exposed to cold temperature (45). We speculated that a similar mechanism operates during *V. parahaemolyticus* biofilm formation.

The bacterial T3SS nanomachine, delivering effector proteins directly from the bacterial cytosol to host cells (46,47). However, several studies concluded that the expression of T3SS genes were repressed in biofilm-growing bacteria (48). Calder et al. showed that the T3SS1 induction has no influence on biofilm growth (34), Ferreira et al. demonstrated that T3SS1 genes expression and biofilm production were inversely regulated (49). In this study, we found that genes encoding T3SS1 have significantly down regulated after cold shock. The results were consisted with the research of Ferreira et al. which indicated the expression of T3SS1 genes can regulate the biofilm formation of *V. parahaemolyticus* (49).

The T3SS mutants impaired the adherence to surfaces during biofilm formation and caused changes in the expression of proteins involved in metabolic processes, energy generation, EPS production and bacterial motility as well as outer membrane proteins (50). Moreover, EPS production and bacterial motility were also altered in the T3SS mutants. After cold shock, the change of EPS production may be induced by the regulation of T3SS genes. Jennings et al. also reported that the T3SS encoded by *Salmonella* Pathogenicity Island 1 (SPI1) can mediate biofilm-like cell aggregation (51). In addition, the flagellum gene and T3SS gene are involved in regulating the biofilm formation at low temperature, which may help *V. parahaemolyticus* to adapt the adverse environment.

In this study, we demonstrated that cold shock induced the expression of related genes of biofilm formation and increased biofilm biomass continuously, causing that the biofilm cells had the ability to adapt to the low temperature shift. It is interesting to investigate how biofilm cells to sense the cold shock signal to regulate the gene expression in further research. More research on the molecular mechanisms between biofilm formation and those related genes is recommended in the future. Besides, an important aspect is to explore the related stress proteins, antifreeze proteins and possible role of the membrane proteins in sensing environmental temperature, such as cold-shock proteins (CSPs) and cold acclimation proteins (Caps). They may be overexpressed during prolong growth of the cold-tolerant bacteria at low temperature.

In conclusion, cold shock (4 °C and 10 °C) could induce the changes of genes expression related with biofilm and impact biofilm structure, causing that the survival biofilm cells adapt to the low temperature shift gradually. Therefore, bacterial pre-formed biofilm exposed to cold shock was not entirely inhibited, which still be a potential risk in nature environmental.

## 4. MATERIALS AND METHODS

### 4.1 Bacterial strains and culture conditions

*V. parahaemolyticus* strain (ATCC17802) was inoculated from storage at - 80 °C into thiosulfate citrate bile salts sucrose agar (TCBS agar, Land Bridge Technology, Beijing, P.R.China) and incubated overnight at 37 °C. Single colony on the TCBS agar was cultured into 9 mL of Tryptic Soy Broth (TSB, Land Bridge Technology, Beijing, P.R.China) containing 3 % (w/v) NaCl and incubated overnight at 37 °C with shaking at 200 rpm. The *V. parahaemolyticus* cultures were diluted with fresh TSB (3 % NaCl) to an OD_600_ value of 0.4 for subsequent experiments.

### 4.2 Biofilm formation assay

Static biofilms were grown in 24-well polystyrene microtiter plates (Sangon Biotech Co., Ltd., Shanghai, P.R.China) and biofilm production was measured using crystal violet staining with some modification (52). In brief, the growth of biofilms was initiated by inoculating 10 μL of the cell suspension which cultured in 2.1 into 990 μL of fresh TSB medium (3 % NaCl) in the individual wells. All samples were statically incubated at 37 °C for 24 h to obtain the pre-formed biofilm, then the pre-formed biofilm was shifted to low temperature (4 °C, 10 °C) for 12 h, 24 h, 36 h, 48 h and 60 h without shaking, respectively. To prevent the medium evaporation, all plates were sealed with plastic bag. At each of these time points, the biofilms were quantified by crystal violet staining. Specifically, the suspension was discarded and the wells were gently washed three times with 1 mL of 0.1 M phosphate buffer (PBS) to remove non-adhered cells, and subsequently the biofilm was stained with 1 mL of 0.1% (w/v) crystal violet (Sangon Biotech Co., Ltd., Shanghai, P.R.China) for 30 min, then washed three times with 1 mL of 0.1 M PBS to remove unbound crystal violet. After drying for 45 min in a 60 °C incubator, biofilm stained by crystal violet was dissolved in 2 mL of 95% ethanol (Sinopharm Chemical Reagent Co., Ltd., Shanghai, P.R.China) for 30 min. Wells containing un-inoculated TSB served as negative controls, and the difference between the optical density of tested strains and negative control (OD_570_) was used to characterize the biofilm-forming ability of the tested strains (53). This experiment was tested in triplicate.

### 4.3 EPS analysis

The exopolysaccharides in *V. parahaemolyticus* biofilm were extracted using sonication method with some modification as described previously (54,55). Biofilm cells on the wells were removed by vortexing and scraping after addition of 1 ml 0.01 M KCl (Sinopharm Chemical Reagent Co., Ltd., Shanghai, P.R.China). The cells were disrupted with a sonicator (JY92-IIN, Ningbo scientz biotechnology Co., Ltd., Ningbo, P.R.China) for 4 cycles of 5 s of operation and 5 s of pause. The sonicated suspension was centrifuged (4,000 g, 20 min, 4 °C), and the supernatant was filtered through a 0.22 μm membrane filter (Sangon Biotech Co., Ltd., Shanghai, P.R.China), 100 μL of the filtrate was mixed with 200 μL 99.9 % sulfuric acid in new tubes. After incubation at room temperature for 30 min, 6 % phenol was added to the mixture. Then after incubation at 90 °C for 5 min, the absorbance of mixture was measured at 490 nm. The amount of carbohydrate was quantified by dividing OD_490_ / OD_595_ values.

The amount of proteins was quantified by the Lowry method (55). 40 μL of the filtrate was mixed with 200 μL Lowry reagent (L3540, Sigma Aldrich, St. Louis, Missouri, USA) in new tubes. After incubation at room temperature for 10 min, 20 μL Folin-Ciocalteu reagent (L3540, Sigma Aldrich, St. Louis, Missouri, USA) was added to the mixture. Then after incubation at room temperature for 30min, absorbance was measured at 750 nm. The amount of proteins was quantified by dividing OD at 750 nm by OD at 595 nm.

### 4.4 Quantitative viable bacteria under biofilm and planktonic conditions

The viable bacteria under biofilm and planktonic conditions were quantified by plate counting method. Biofilm-forming bacteria on the well were removed by vortexing and scraping after addition of 1 ml 0.85% NaCl. Planktonic *V. parahaemolyticus* was cultured as described in 2.1, and all samples were incubated at 37 °C for 24 h, then they were shifted to low temperature (4 °C or 10 °C) for 12 h, 24 h, 36 h, 48 h and 60 h, respectively. Enumeration of *V. parahaemolyticus* was performed by serial dilution and spreading onto TCBS agar plates. The plates were incubated at 37° C for 24 h and colonies were counted.

### 4.5 Confocal laser scanning microscopy imaging

Quantitative parameters describing biofilm physical structure have been extracted from three-dimensional CLSM images and used to compare biofilm structures, monitor biofilm development, and quantify environmental factors affecting biofilm structure. *V. parahaemolyticus* biofilm was observed using a confocal laser scanning microscopy (CLSM). The biofilm was immobilized using 2 mL of 4 % (v/v) glutaraldehyde solution for 2 h at 4 °C, and rinsed 3 times with 0.1 M PBS and stained with SYBR Green I (Sangon Biotech Co., Ltd., Shanghai, P.R.China) for 30 min in darkness at room temperature (56). The wells were then washed with 0.1 M PBS to remove the excess stain and air dried. CLSM images were acquired using a Zeiss LSM710-NLO Confocal Laser Scanning Microscopy (Carl Zeiss, Jena, Germany) with a 20× objective. Biofilm structure morphology in three dimensions was analyzed using Image Structure Analyzer-2 (ISA-2) (57,58).

### 4.6 RNA extraction, reverse transcription, and RT-PCR analysis

Total RNA from biofilms was extracted and purified using the RNA extraction kit (Generay Biotech Co., Ltd., Shanghai, P.R.China), according to the manufacturer’s instructions. RNA concentrations were determined by measuring the absorbance at 260 nm and 280 nm (ND-1000 spectrophotometer, NanoDrop Technologies, Wilmington, DE, USA). Reverse transcription (RT) was performed with 200 ng total RNA using the Prime Script RT reagent Kit with gDNA Eraser (Takara, Dalian, P.R.China) following the manufacturer’s instructions.

The qPCR reaction mixture (20 μl) contained 10 μl mix, 0.4 μL ROXII, 0.8 μM of the appropriate forward and reverse PCR primers, 2 μl of template cDNA and ddH_2_O 6μL. The reactions were preformed using an Applied Biosystems 7500 Fast Real-Time PCR System (Applied Biosystems, Carlsbad, USA). Negative controls (deionized water) were included in each run. Amplifications were performed in duplicate. The expression levels of all of the tested genes were normalized using the 16S rRNA gene as an internal standard (59). Relative quantification was measured using the 2^−ΔΔCt^ method (the amount of target, normalized to an endogenous control and relative to a calibrator, where ΔΔCt = (Ct target − Ct reference) sample − (Ct target − Ct reference) calibrator) (60). The specific primers (Table1) were designed by Primer Premier 5.0 software, and all synthesized by Shanghai Sangon Biotech Company.

### 4.7 Statistical analysis

The statistical analysis was performed using the SPSS statistical software (version 17.0; SPSS Inc., Chicago, IL, USA), including two-way Analysis of Variance (ANOVA) for time-course evaluations, the Student *t*-test for comparison between groups, Pearson correlation coefficient at the 0.01 and 0.05 significant level. Values were considered significantly different if *p* < 0.05. Calculations and figures were performed using Microsoft Excel 2007 (Microsoft Corporation, Redmond, WA, USA) and origin 8.0, respectively. Linear regression analysis using Excel 2007.

## ACKNOWLEDGMENTS

This research was supported by the National Natural Science Foundation of China (31571917 and 31671779), Shanghai Agriculture Applied Technology Development Program (Grant No. G20150408, G20160101 and T20170404), Innovation Program of Shanghai Municipal Education Commission (2017-01-07-00-10-E00056), the “Dawn” Program of Shanghai Education Commission (15SG48).

